# Genome build information is an essential part of genomic track files

**DOI:** 10.1101/120923

**Authors:** Chakravarthi Kanduri, Diana Domanska, Eivind Hovig, Geir Kjetil Sandve

## Abstract

Genomic locations are represented as coordinates on a specific genome build version, but the build information is frequently missing when coordinates are provided. It is essential to correctly interpret and analyse the genomic intervals contained in genomic track files. Here, we demonstrate that this crucial metadatum (or rather datum) is often isolated from the genomic track files in public repositories and journal articles, which could be a major time thief. We propose best practices to ensure that genome build version is always carried along with genomic track files. Although not a substitute to the best practices, we also provide a tool to predict the genome build version of genomic track files.

## Background

The data deluge that arose with the advent of high-throughput sequencing methods needs no introduction [1]. To enable reproducibility and reuse of the data, a community-driven common practice encourages researchers to deposit the generated data to public repositories [2,3]. Larger research consortia that have generated data spanning billions of base pairs across a plethora of individuals, cell types and experimental conditions, have also made their data public [4–6]. This good practice facilitates the reuse of the data to dig up further biological insights (e.g., see [7,8]). However, the reuse of data in such manners depends largely on the availability of metadata, which amongst other information describes the essential experimental and data processing details associated with the dataset [9,10]. Still, we find that sufficient metadata is often lacking for datasets in both public repositories and journal articles.

One fundamental, but surprisingly common missing information element of genomic datasets is the genome build version that a dataset relates to. This is especially problematic for file formats solely containing genomic intervals. Just as the start and end positions (coordinates) in a BED- or GFF-file do not provide any information without the knowledge of which chromosome the positions (offset) refers to, the chromosome and position together (genomic coordinates) also do not denote any meaningful location without the version of a genome build. In other words, without a genome build information, the sequence coordinates would just be house numbers without a street name. Therefore, for files that exclusively provide data in the form of coordinates on a reference genome, such as BED, WIG or GFF, the genome build version is not only critical, but even a part of the data itself (it is actually data, not metadata).

Although the failure to supply the genome build information often occurs, also the way this vital information is being collected and stored is an equally big concern. Several of the public repositories and journals recommend the submission of a range of metadata (including genome build information) [9,10], which are usually stored and provided in a separate file or webpage (but not as an integral part of the data file itself). In other words, the common file formats that solely contain genomic intervals do not necessarily carry genome build information in any form (e.g., **Table 1**). Owing to the largely collaborative nature of genomics research (as of now), the data may travel between several computers back and forth during a project, thus requiring the documentation and explicit specification of genome build information every single time (e.g., in email or otherwise). However, this process is error-prone and is prone to failure. Eventually, lack of genome build information could become a major time thief that may also potentially lead to erroneous data integrations. Here, we demonstrate the frequent isolation of genome build information from genomic track files and propose that this information should rather be an essential part of the data file itself.

**Table 1:**
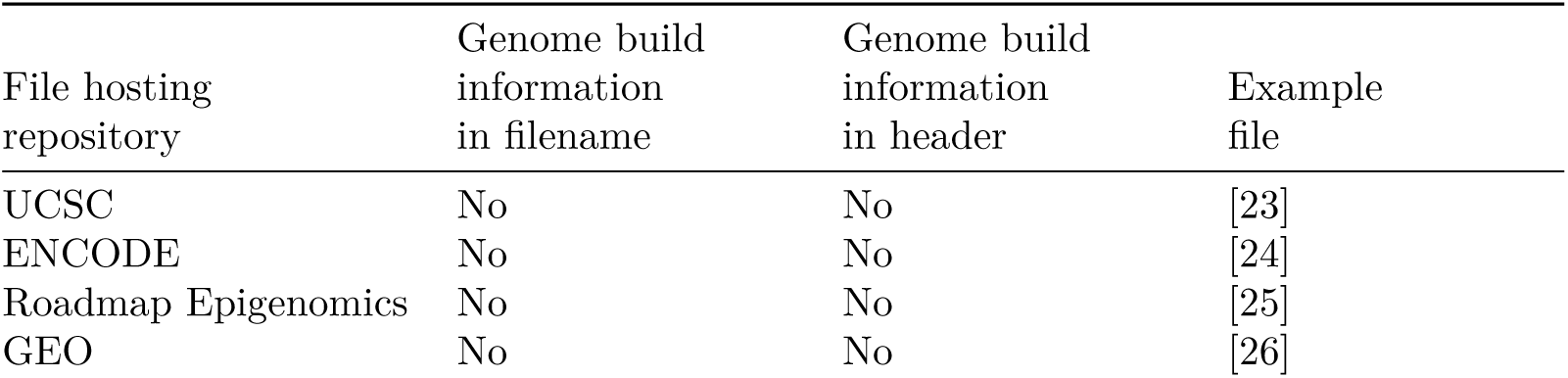
Examples of detachment of genome build information for files downloaded from public repositories.

## Methods and Results

### Extent of incompatibility between two genome-build versions

One of the common consequences of missing genome build information is the integration of genomic coordinates from two different genome build versions, which could largely be erroneous. To exemplify this error, we checked the extent of compatibility between two versions of human genome builds, hg19 and hg38. For this, we downloaded the size of both autosomes and sex chromosomes for hg19 from the UCSC database using fetchChromSizes [11] and segmented the chromosomes into 1 kb bins, resulting in a total of 3095689 bins. The genomic coordinates of the bins were then lifted over to hg38 using the UCSC liftover tool, where the conversion failed for 238542 bins (7.7% of total bins). For the bins that were common between both hg19 and hg38, we computed the distance between the midpoint of each bin on hg19 and its corresponding bin (after lift-over coordinates) on hg38 to know whether the genomic coordinates remained identical between genome builds. This revealed that only ~ 18696 bins (0.65 % of the total number of bins) had identical genomic coordinates on both hg19 and hg38 for autosomes and sex chromosomes, and ~ 89.0% of the total bins were further apart than 30 kb. Overall, integration of genomic intervals between these two builds would be erroneous for ~ 99.4% of total bins (3076993000 / 3095689000 bases). Despite the large discrepancy, this problem could easily be overlooked when performing genome arithmetic operations because of the common sequence names and coordinates between different versions of genome builds. For example, see **Figure 1**.

**Figure 1:**
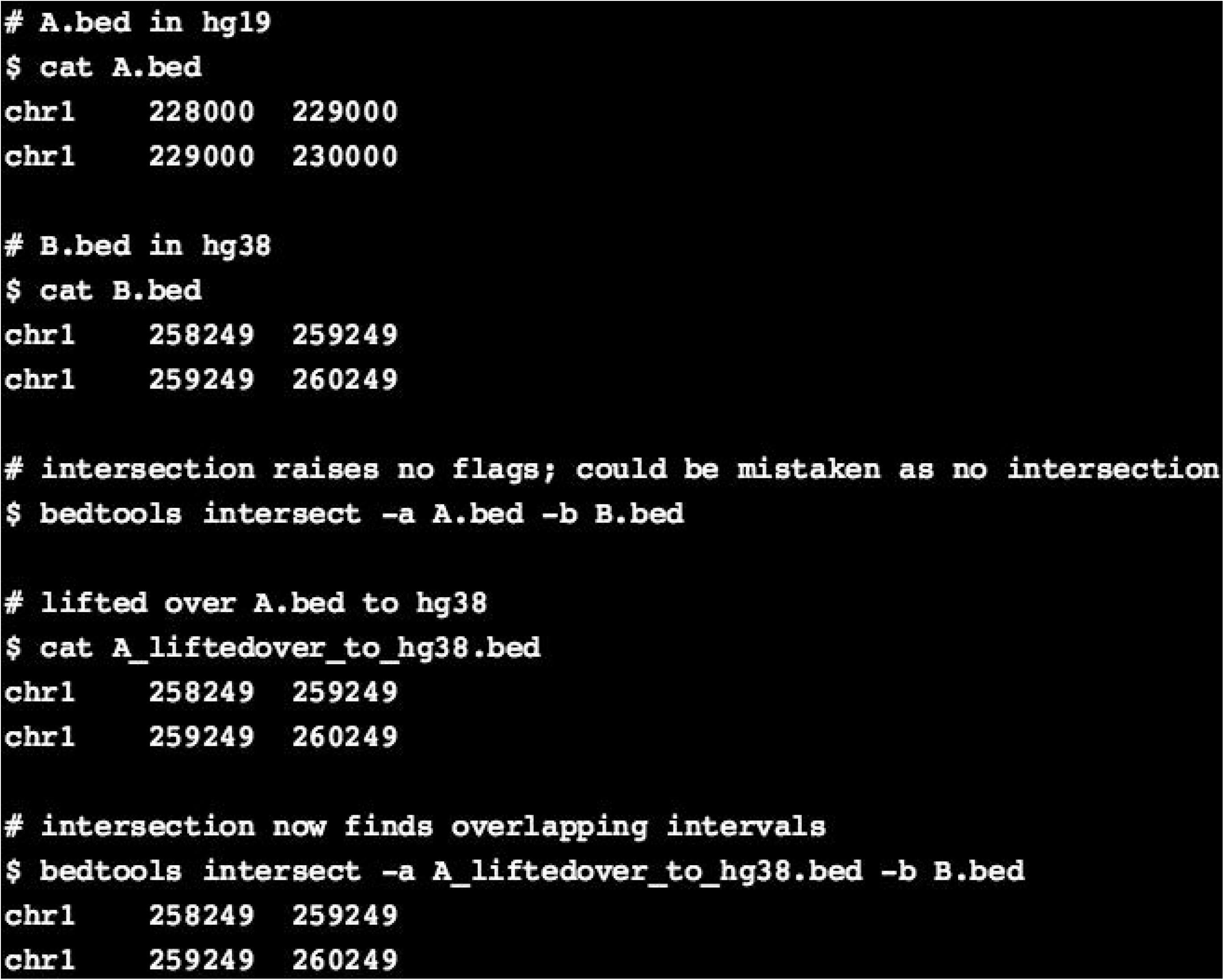
Example of erroneous genome arithmetic operations because of integrating data from incompatible genome builds. The sequence intervals in A.bed (hg19) correspond to the intervals in B.bed (hg38). However, when genome arithmetic operations (like intersection) are performed in the absence of genome build information, there will be no overlap between the intervals in both the files, as the coordinates appear non-overlapping. If it is known that the files correspond to two different genome builds, lift-over of coordinates in one file, would result in the detection of overlapping genomic intervals between both the files.

### Genome build information in a sample of repositories and journals

We first determined whether genome build information is consistently supplied along with submissions to public repositories. As a representative example, we examined the records in the GEO and ENCODE databases with the following search criteria: In the GEO database, we examined all the records (one sample per series) that involved high-throughput sequencing submitted after 31.12.2008 for three species: *Homo sapiens, Mus musculus* and *Drosophila melanogaster*. We then checked whether the data processing section of metadata explicitly mentioned the genome build information, by case-insensitively searching for the following words: {hg17|hg18|hg19|hg38|grch36|grch37|grch38|build37.2|build37.1|build36.3|ncbi35|ncbi36|ncbi37|mm8|mm9|mm10|grcm38|bdgp6|bdgp5|bdgp5.25|build5.41|build5.3|build5|build4.1|dm6|dm3|ncbi}. In the ENCODE database, we examined the metadata file of all records.

Around 23.0% of the queried series records did not contain the genome build information explicitly in the data processing section of metadata in the GEO database (queried on 17.03.2017 according to the search criteria stated above), whereas all the relevant records in the ENCODE database contained genome build information in the metadata section.

Next, using a similar set of search criteria, we retrieved a total of 6155 articles across four journals. We then employed a series of filtering steps to shortlist articles that had a GEO accession ID corresponding to a high-throughput sequencing experiments, resulting in 332 articles. Of those, ~ 14.0% of the articles did not mention genome build information in the full text of the article, but it was mentioned in the metadata section of the GEO database. On the other hand, ~ 16.0% of the articles mentioned the genome build information in the full text of the article, but not in the GEO database, from where the data is usually downloaded by other researchers. ~ 4.0% of the articles did not supply genome build information to neither journal nor repository (detailed in additional material; **Additional Tables 1-2**).

### Detachment of genome build information from genomic track files

To demonstrate the extent of detachment of genome build information from data files, we next downloaded several files from public repositories and checked whether genome build information was carried along with the files in some form. First, we checked for the file formats BED and GFF, that are attached as supplementary file to sample records in the GEO database. Again, after restricting the search criteria to three species: *Homo sapiens, Mus musculus* and *Drosophila melanogaster*, we retrieved 967 BED files and 2100 GFF files. We then checked whether the filenames or the header lines (we checked the first 50 lines) explicitly mentioned the genome build information. For this, we again searched for the names of specific genome builds of the three species listed above. Overall, while ~ 46.0% of the total BED files from the GEO database carried genome build information, only ~ 0.6% of the total GFF files carried it either in filename or in header (**Table 2**).

**Table 2:** Genome build information carried in filenames or header lines.

| Files | Total checked | Only in filename | Only in the file | In both |
| --- | --- | --- | --- | --- |
| GEO: BED | 967 | 149 (15.4%) | 213 (22%) | 84 (8.7%) |
| GEO: GFF | 2105 | 0 | 0 | 12 (0.6%) |
| ENCODE: BED | 26,503 | 0 | 0 | 0 |

Further, we downloaded 26503 BED narrowPeak files (from a total of 4775 records: *Homo sapiens* - 3533, *Mus musculus* - 994 and *Drosophila melanogaster* - 248) from the ENCODE database, and repeated the same analysis. We found that none of the files carried genome build information in either their filename or as part of their header. Although these statistics (**Table 2**) largely stem from the fact that these file formats do not require genome build information as an obligatory field, this exemplifies the extent to which genome build information is detached from the genomic interval files after downloading them from public repositories. Once such genomic interval files are downloaded and exchanged between computers, one cannot totally exclude the possibility of failure to recording the genome build information, thereby leading to additional time investment in procuring the genome build information, or resulting in erroneous integrations.

### Examples of attaching genome build information to genomic track files

The examined records that did store genome build information, stored it in any of the following ways: (a) a header line (commented line) that specified genome build version (e.g. [12]), (b) as part of the ‘track name’, which is usually the header line that appears in the track files of genome browsers (e.g. [12]), (c) recorded the path of input files in the header, where the path contained genome build information (e.g. [13]), (d) made use of required fields, like ‘source’ or ‘feature’, to include genome build information (e.g. [14]), or (e) as part of the filenames [14]. Further, we found that customized file formats in some instances dedicated a column to store genome build version (e.g. [15]).

### Recommendation on ways to specify genome build

To ensure that genome build information stays with the data, we recommend that the information is included as part of initial header/comment lines inside the file itself. For most file formats, a comment line starting with hash (#) would allow the information to be human readable, without disturbing parsers. This is explicitly supported e.g. in file formats like GFF, [16] and also appear to be allowed *de facto* in formats like BED [17] (though not formally specified as part of the format [18]). For BED files, as an alternative, one could add “genome=xxx” as part of a “track”-prefixed header that is allowed in the track files used by genome browsers [18]. In cases where adding information to the data contents of a file is not possible/practical, an alternative is to specify genome build as part of the file name. Although the file name may change as it moves between people and computers, it is often stable. Having the genome build only as part of the full path/URL of a file is risky, as it will be disrupted by almost any transfer. A last option is to simply specify the genome build for every data line along with the sequence (chromosome) identifier, although superfluous for datasets where all regions are from the same genome build.

### Tool to predict the genome build version

Although not a substitute to explicitly supplementing the genome build information, we here provide a tool to predict the genome build version of orphan genomic track files. The tool is available at [19] on the Genomic HyperBrowser [20], an integrated open-source tool for statistical genome analysis. The web tool accepts a wide-range of genomic track file formats and currently supports 19 species. In addition, we provide a command-line tool as an R package that supports human, mouse and Drosophila genome builds [21]. When we tested the webtool on public data from ENCODE, the tool predicted the correct genome build for 98.2% of the broad peak files (n=223) for the K562 cell line. We noticed that the tool failed when the genomic track files did not strictly adhere to the file format specifications (see Additional material). Notably, any such tool cannot distinguish between genome builds if the genomic track files of interest do not contain sequence coordinates that are unique to a genome build version. In other words, the prediction is infeasible if all the input sequence coordinates are equally compatible with two or multiple genome builds.

## Conclusion

This study demonstrates the detachment of genome build information from genomic track files in both public repositories and journal articles, which could lead to additional time investment in inferring the genome build or potential erroneous genome arithmetic operations. The findings also exemplify the extent of incompatibility between the sequence coordinates of two genome build versions that would result in erroneous integrations when performing genome arithmetic operations. We propose three ways to ensure that genome build information is always carried along with genomic track files, where the preferable solution is to record it as part of the header lines in genomic track files. To facilitate the adoption of orphan genomic track files, we provide a tool that predicts the genome build version.

## Declarations

## Ethics approval and consent to participate

Not applicable

## Consent for publication

Not applicable

## Availability of data and material

All the scripts used for web crawling, querying the databases, retrieving the files and all URLs of the files retrieved are submitted to Zenodo database and can be accessed at [22].

## Authors’ contributions

CK and DD performed the database queries, web crawling, analyses, and tool implementations. EH conceived part of the analyses and tool implementation. GKS conceived the idea and part of the analyses. CK and GKS drafted the manuscript. All authors read and approved the final manuscript.

## Acknowledgements

We thank Sveinung Gundersen and Lex Nederbragt for helpful discussion.

